# What drives male association tendencies in wild zebrafish? Role of female and vegetation densities

**DOI:** 10.1101/2020.01.14.907139

**Authors:** Aditya Ghoshal, Anuradha Bhat

## Abstract

Mating strategies in species is context-dependent and driven by several ecological and demographic factors. In natural habitats, a multitude of ecological factors interact and these eventually determine mating preferences and mate choice decisions among species. While zebrafish (*Danio rerio*) is a widely studied biological model organism, our understanding of their mating preferences, strategies and their underlying ecological drivers are still limited. Here, we explore the role of ecological factors such as spatial variation (or patchiness) in mate density and associated vegetation cover in determining mate association tactics among males in a wild zebrafish population. We employed a multi-choice experimental design for a better representation of ecologically relevant scenarios. Our results revealed that when presented with patches with increasing female densities, males displayed only a marginal increase in preference for higher female density patches. However, when female density varied concomitantly with variation in vegetation cover, the males associated more with higher foliage density patches irrespective of the female density in that patch. Our findings throw light on the complex interaction between these two most basic ecological factors in determining mate search strategies and mate associations among these group-living fish species.

## INTRODUCTION

Mating decisions among species are the result of a trade-off between benefits from mating events and the combined costs of gamete production, mate search and mate sampling (Wagner, 1998). Mate choice decisions would depend on several factors such as density of potential mates and their encounter rates, presence of other competing individuals (Weiss and Dubin 20182) or presence of predation threat (Forsgren 1992; Bierbach et al. 2011), along with innate preferences for any phenotypic trait in a prospective mate (Willis et al. 2011). Trivers (1972) considered females as the primary driver of sexual selection in several animals, due to their higher cost of egg production (relative to sperms) and post-fertilization parental care. However, in a natural environment, mate choice may not only depend on preferences of the females but also that of males. For example, in species with sex-role reversals and male parental care we do see males being the choosy sex (Eens and Pinxten 2000; Gwynne 2016). Male mate choice can, however, evolve even in absence of male parental care, under specific ecological contexts (Edward and Chapman 2011). Mate search is not only time-expensive but also an energy-expensive process. Here, we investigated mate association and decision-making in male zebrafish and the role of specific ecological factors in determining their association preferences.

Among fish species, mate choice is determined by a combination of factors such as mate quality, mate density, presence of competitors, etc. (Dugatkin and Fitzgerald 1997). Mate search by males is not only time-expensive but also an energy-expensive process. Mate search can also put the males under increased predation risk (Forsgren 1992; Berglund 1993). Population density (Jirotkul 1999) and operational sex ratios (Kvarnemo and Ahnesjo 1996; Weir et al. 2011) also determine the search tactics. In the face of all these costs, it is important for the male to maximize its mating probability based on the variation in number of available receptive females along with constantly changing ecological factors.

Zebrafish is one of the most popular model systems in biological studies that are bred worldwide in laboratory environments (Nasiadka and Clark 2012). While there have been detailed explorations on their mating and courtship behavior (Darrow and Harris 2004), we still understand little of their mating strategies and the ecological factors that influence them. Mating studies in zebrafish have generally focused on studying female preference for phenotypic traits such as body size (Pyron 2003), novel phenotypes (Owen et al. 2012) and dominance (Spence and Smith 2006) along with egg allocation strategies in female (Uusi-Heikkilä et al. 2012; Skinner and Watt 2007). However, there still exists a large gap in our current understanding of the mating strategies in this species. In nature, they are believed to be group spawners, with females scattering their eggs in specific oviposition sites (Spence et al. 2007). Zebrafish hatchlings are born as females and switch over later in life to males due to factors that are still partially understood. Zebrafish is phenotypically monomorphic with no apparent morphologically visible sexual traits or display colorations. Females are visually distinguished by the curvature of the belly when they are gravid. They do not show parental care and both sexes show egg cannibalism (reviewed in Spence et al. 2007). Studies on mating ecology of species have focused on animals with large ornaments or phenotypic traits which have allowed easier testing of hypotheses on mate selection and mating strategies. The lack of sexual traits in zebrafish can be a hindrance for such traditional approaches to studying mating dynamics. However, it also provides us with the opportunity to expand our knowledge of mating ecology in such sexually monomorphic species due to the wide usage of zebrafish in biological research and well-developed genomic tools to understand the physiological basis of their mating ecology.

In nature, males could encounter females in varying numbers and this would allow males to choose their association with specific potential female partners. Here, we focused on whether male mate search tactic is influenced by the number of gravid females aggregated spatially in patches and how associated vegetation cover can modulate the search tactics. In contrast to commonly used two-choice tests, we used a multi-choice test paradigm to test the males’ preferences. We expected male association to greater in patches with higher female densities.

Presence of vegetation cover in aquatic habitats is an important ecological feature that is known to regulate foraging behavior of predators (for example, in Spotted gar *Lepisosteus oculatus*, Ostrand et al. 2004) and also provide protection and shelter to prey species (Christensen and Persson 1993). Indeed, zebrafish are known to prefer enriched vegetated areas compared to the bare tank (Delaney et al. 2002; Schroeder et al. 2014). Carfagnini et al. (2009) found that female zebrafish showed reduced aggression in spatially complex habitat compared to a bare habitat. However, the role of vegetation cover in mate search tactics is largely unclear. In this study, we also investigated whether male association preferences are influenced by varying vegetation densities. Specifically, our study addressed the following questions:

a. Does varying number of receptive females influence the preference/choice of a patch by the males and thus influencing the association pattern?
b. Does vegetation density associated with female patches (varying in number) change the association pattern?

We expected the males’ association to a patch would be positively related to the number of females present in that patch indicating a strategy of optimization of resources by choosing the patches with a greater probability of finding a suitable mate. We also expected that in presence of vegetation they would prefer patches with higher degree of vegetation cover associated with a higher number of females.

## MATERIALS AND METHODS

### Procuring subject animals and maintenance

We used wild-caught zebrafish (from Howrah district, West Bengal, India), bought from a commercial supplier. The fish were maintained in the laboratory in mixed-sex groups of approximately 60 individuals in well-aerated holding tanks (60×30×30 cm) filled with filtered water. The lighting in the laboratory was maintained at 14hL:10hD to mimic the natural LD cycle in zebrafish. They were fed commercially purchased freeze-dried blood worms once a day alternating with brine shrimp *Artemia*. The holding tanks were provided with standard corner filters for circulation. They were maintained in the laboratory for six months before experiments were conducted to ensure they were all adults and were reproductively mature. Holding room temperature was maintained between 23°C-25°C.

### Experimental setup

The experiments were conducted in a square glass arena (83X 83 cm), with a half-diagonal of the square from the center that approximated ten fish standard body lengths (i.e. 40cm, assuming one body length of adult zebrafish to be about 4cm) (Figure 1). Each corner of the arena was provided with a square chamber (of sides 10 cm) built from transparent mesh for housing the females. This design allowed for the stimuli females to be localized in the patches and not escape into the arena while simultaneously ensuring that the test males can have visuo-chemical communication with the females. The center of the arena was provided with a removable chamber (with holes) for acclimation of the males prior to the trial. Three sets of experiments were performed to test their association preferences under 1) only varying female densities 2) increasing female and vegetation densities and 3) increasing female densities with decreasing vegetation.

**Fig 1.**
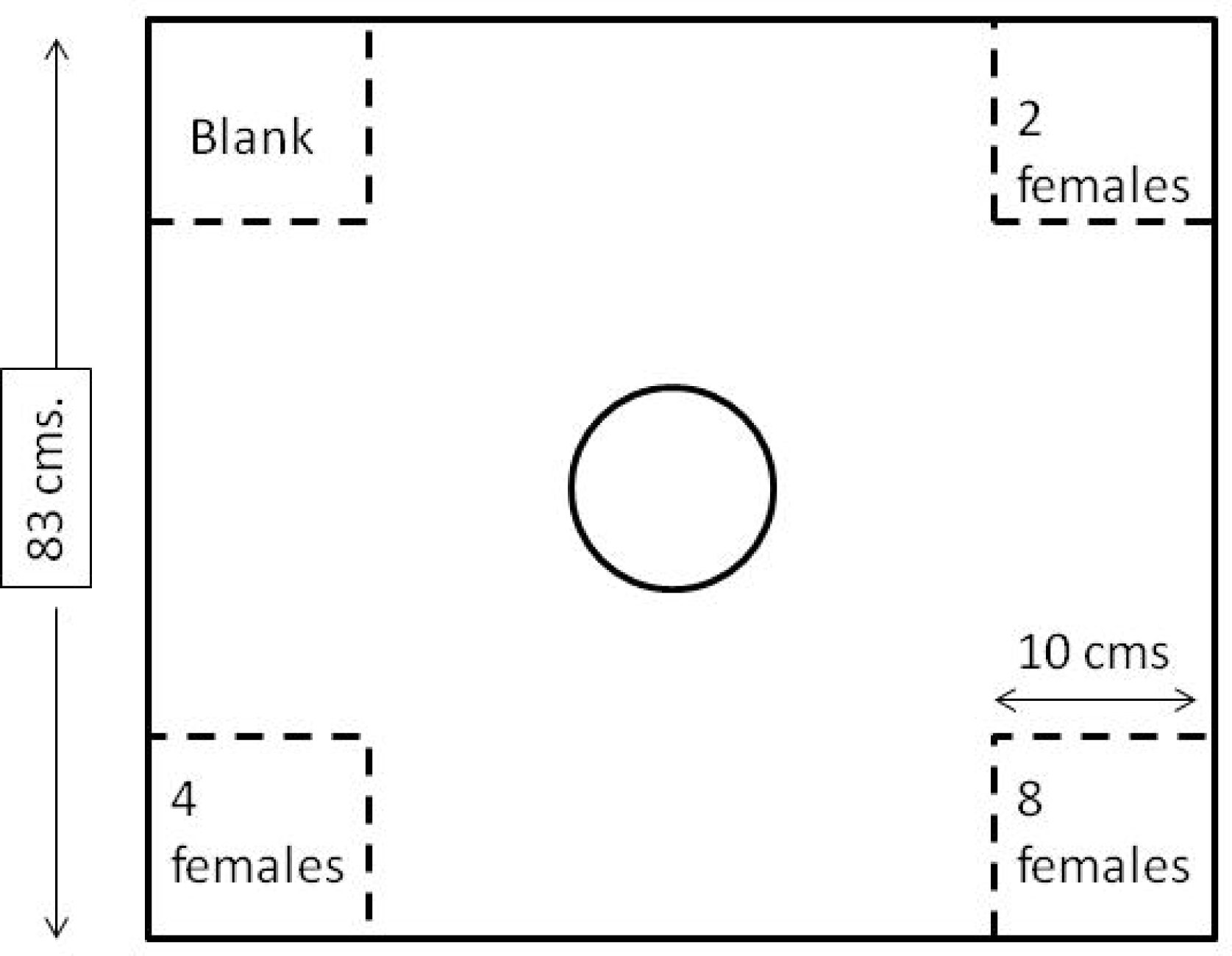
Diagrammatic representation of the arena for the density experimental set-up. The central chamber (indicated by a circle) represents the area where the test males were released and the corner square chamber (separated by transparent mesh) contained females of varying density. The distance of each patch from the central chamber was 40 cms.

### Association preference experiment with varying female densities

For this experiment, each small chamber within the arena housed two (low number), four (medium number), eight (high number) or no (blank) females. These chambers represented patches of varying female numbers. The position of the females, as well as the composition of females within each patch, were randomized between trials. A total of 20 males were tested for their association preferences.

### Association preference experiment with vegetation

For this experiment, the female-housing chambers (patches) were provided with vegetation cover (using artificial plants) of varying cover density (Fig. 2). Each subject fish was tested under two experimental settings. In E1, the number of females was proportional to the density of associated vegetation cover. We used four different densities of females, each associated with different densities of plants—i) one female + no plants (no cover -N), ii) two females + two plants (low cover - L), iii) four females + three plants (moderate cover -M) and iv) eight females + five plants (high cover -H). For E2, patches with increasing female densities were provided with a corresponding decreasing vegetation cover, except for the patch with no plant cover. That is, there was an interchange in the vegetation cover for the two and eight female patches- i) one female + no plants (no cover -- N), ii) two females + eight plants (high cover -- H), iii) four females + three plants (moderate cover -- M) and iv) eight females + two plants (low cover -- L). All test males were tested in E1 and E2 on consecutive days in no particular order

**Fig 2.**
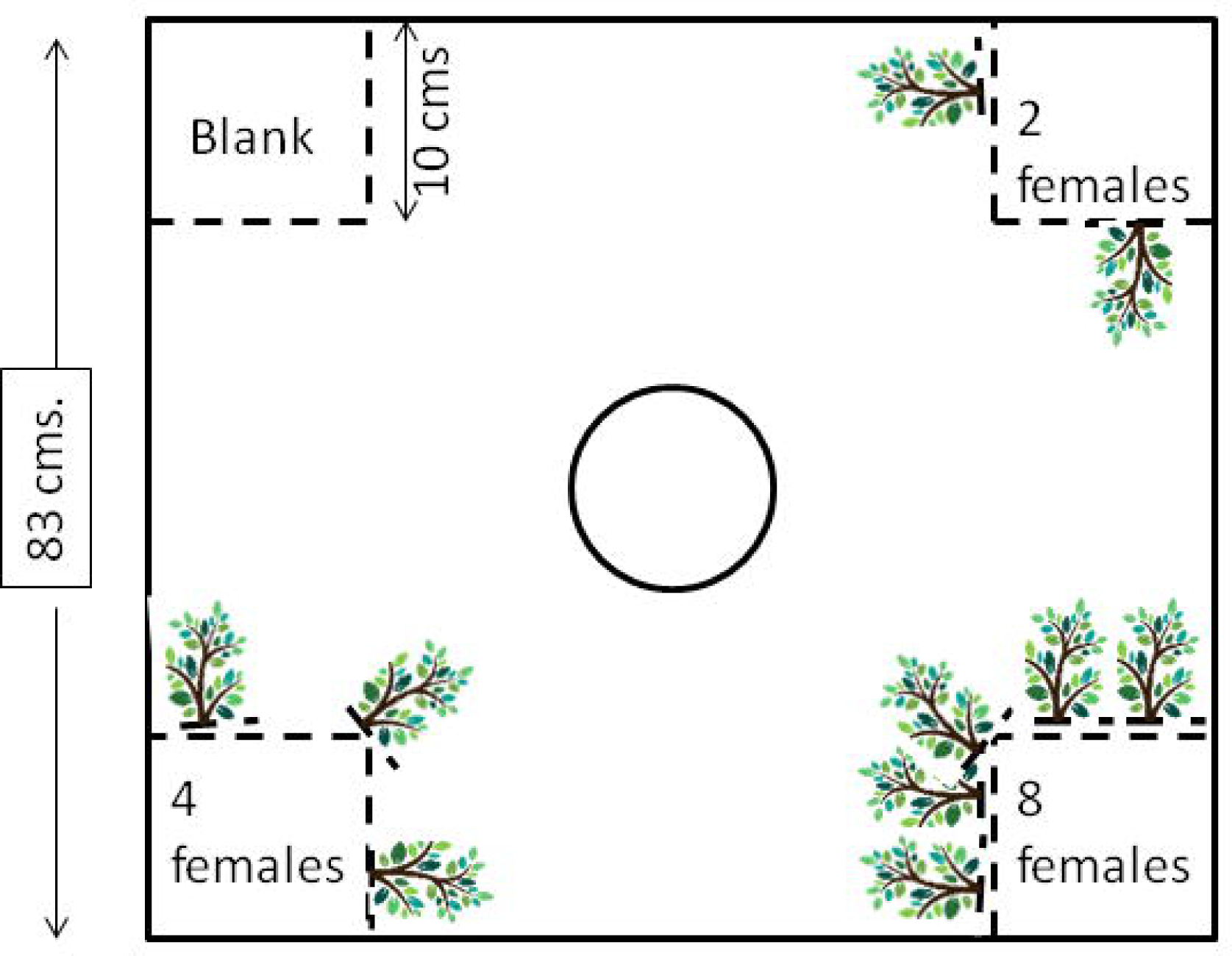
Diagrammatic representation of the arena the vegetation experimental set-up. The central chamber (indicated by a circle) represents the area where the test males were released and the corner square chambers (separated by transparent mesh) contained females of varying density and each patch was associated with variable number of plastic trees representing vegetation cover.

### Experimental Protocol

For the experiment involving association preferences with only varying female numbers a total of 20 males were tested, while 24 males were tested for experiments on the association preferences in varying female numbers combined with vegetation density gradients (E1 and E2 experiments). We isolated subject males of comparable sizes and kept them in individual isolation in 500 ml jars for four days prior to experiments as that allowed us to keep track of individual fish and also stimulated mate-seeking behavior (Roy and Bhat 2015; Gerlach 2006). They were fed freeze-dried blood worms every day at constantly maintained feeding times. The gravid females that were used for the experiment as stimuli for association were isolated (about 22 females) in a small holding tank (30×20×20 cm) with a feeding regimen similar to the test males. Before the start of each trial, we introduced the females into each chamber (patch) randomly (according to the experimental setup described above) and left them there for 15 min. for acclimation. A single male individual was then introduced into the central cylindrical chamber open at both ends (made of transparent plastic and provided with holes). After a five-minute acclimation period, the chamber was gently removed to allow the male to swim freely in the arena and video recording was commenced. Video recordings were done using a camera (Sony DCR-PJ5, Sony DCR-SX22) placed perpendicularly above the arena. The test fish (males and females) were fed only after the end of experimental trials, on each day of experiments. At the end of the trials, the fish were returned to their holding tanks.

We recorded the behavior of each test fish for a duration of 10 minutes. All videos were analyzed using the software BORIS (Friard and Gamba 2016). A single visit to any of the patch was denoted when the male approaches within 6 cm. (1.5 times their average body length) of the patch. We collected data on three parameters: total number of visits to each patch, the total amount of time spent in each patch and the mean time spent per visit within each patch. The same overall protocol was followed for all sets of experiments.

### Statistical Analyses

We noted the total number of visits to each patch, the total duration of time spent in each patch and mean time spent per visit per patch for the entire ten minutes duration of video recording for each test male.

All statistical analyses were performed in R studio (version 1.1.463). We developed generalized linear mixed models (GLMMs) using package ‘lme4’ (version 1.1-19; Bates et al. 2015) with ‘fish ID’ as the random factor and ‘Patches’ as the fixed factor, with four levels representing the four choices for the test (male) fish. Wherever required, post hoc Tukey’s tests, based on R package ‘multcomp’ (version 1.4-10; Hothorn et al. 2008) or paired Wilcoxon test was performed to compare measurements of the parameters (i.e., total number of visits, total duration of time spent, mean time spent per visit) between the patches.

For analyzing the data for the second and third experiments involving varying female densities along with vegetation densities (E1 and E2), we followed a similar procedure of constructing a GLMM followed by post hoc tests. GLMM models were constructed with a single independent variable “patch” that had four levels designated as H (high tree density), M (moderate tree density), L (low tree density) and N (no tree). We performed post hoc paired Wilcoxon signed rank tests to compare between the patches.

## RESULTS

### Association preference experiment with varying female densities

Data for total number of visits was found to fit negative binomial distribution model (AIC=588.35) and hence we used ‘glmer.nb’ class of glmm in package ‘lme4’. The selected prediction model revealed that the fixed factor,‘patch’ significantly affects the number of visits (AIC=563.9, Wald type II χ2=26.44, df=3, p<0.001) (Table 1a). Post hoc Tukey’s analyses to compare each pair of patches showed that there was a significantly lower number of visits to the patch with no females (blank patch) compared to all three patches—two-female patch (n=20, z=3.5, p=0.002), four-female patch (n=20, z=3.31, p=0.005), eight-female patch (n=20, z=5.06, p<0.001) (Fig. 3a). The data on total duration best fit the log-normal distribution (AIC=831.22) and we used glmmPQL method in ‘MASS ‘package to construct the model. The model showed a significant effect of patch on total duration of time spent (Wald type II χ2=9.36, df=3, p=0.02) (Fig. 3b) (Table 1b). Post hoc Tukey’s test showed significant difference in total time spent between eight-female patch and no-female patch (z=2.81, p=0.02) (Fig. 3b). The mean time spent per visit at each patch was also found to best fit log-normal distribution (AIC=402.73) and glmmPQL method was used to construct the model. Patch was again found to significantly affect mean time spent per visit (Wald type II χ2=10.45, df=3, p=0.02) (Fig.3c) (Table 1c). Post hoc analyses showed there are significant differences in mean time per visit between four-fish patch with the zero-fish patch (z=2.82, p=0.02).

**Table 1.** Summary of results of generalized linear mixed models indicating effect of patch (female number) (fixed factor) and fish id (random factor) on a) total number of visits b) Total duration, c) Mean duration of time spent at each patch. The AIC values of selected models along with estimate values, t-scores, degrees of freedom (df) and p values for each factor are shown. p values≤0.05 is significant. Patch2F, Patch4F, and Patch8F signify patches with 2, 4, and 8 females, respectively.

| Number of visits ~ Patch+(1\|fish ID) |  |  |  |  |
| --- | --- | --- | --- | --- |
| AIC |  |  |  |  |
| 563.9 |  |  |  |  |
| variable | estimate | z-score | df | p |
| Patch2F | 0.51 | 3.5 | 3 | 0.005 |
| Patch4F | 0.48 | 3.31 | 3 | 0.0009 |
| Patch8F | 0.73 | 5.06 | 3 | $<<0.01$ |

b)
| Total duration ~ Patch+(1\|fish ID) |  |  |  |  |
| --- | --- | --- | --- | --- |
| variable | estimate | t-score | df | p |
| Patch2Y | 0.85 | 1.92 | 1 | 0.06 |
| Patch4F | 0.89 | 2.02 |  | 0.05 |
| Patch8F | 1.17 | 2.74 |  | 0.008 |

c)
| Mean duration of time ~ Patch+(1\|fish ID) |  |  |  |  |
| --- | --- | --- | --- | --- |
| variable | estimate | z-score | df | p |
| Patch2F | 0.22 | 0.95 | 1 | 0.35 |
| Patch4F | 0.57 | 2.75 |  | 0.008 |
| Patch8F | 0.5 | 2.34 |  | 0.02 |

**Fig 3a.**
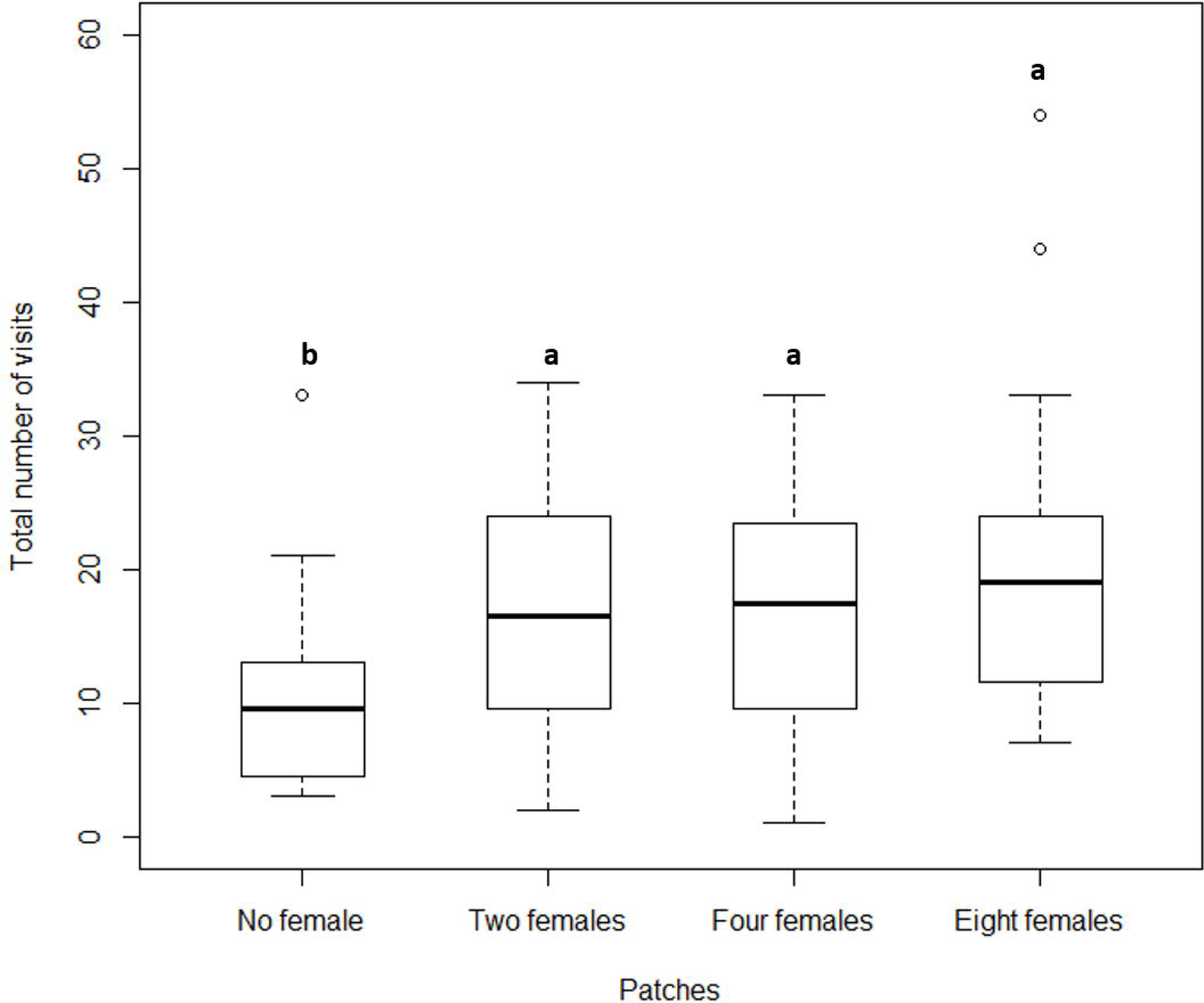
Boxplots showing the total number of visits by the male to the various chambers (patches) containing females in varying density. The chamber containing no females received a significantly lesser number of visits compared to the other three chambers. Differing alphabets indicate statistical significance, while similar alphabets indicate no statistical difference (p<0.05). The lower and upper ends of the box plots represent the 1st and 3rd quartiles, the horizontal line within each boxplot is the median, and the upper and lower whiskers are the 1.5× the interquartile range. Outlines are shown as open circles.

**Fig 3b.**
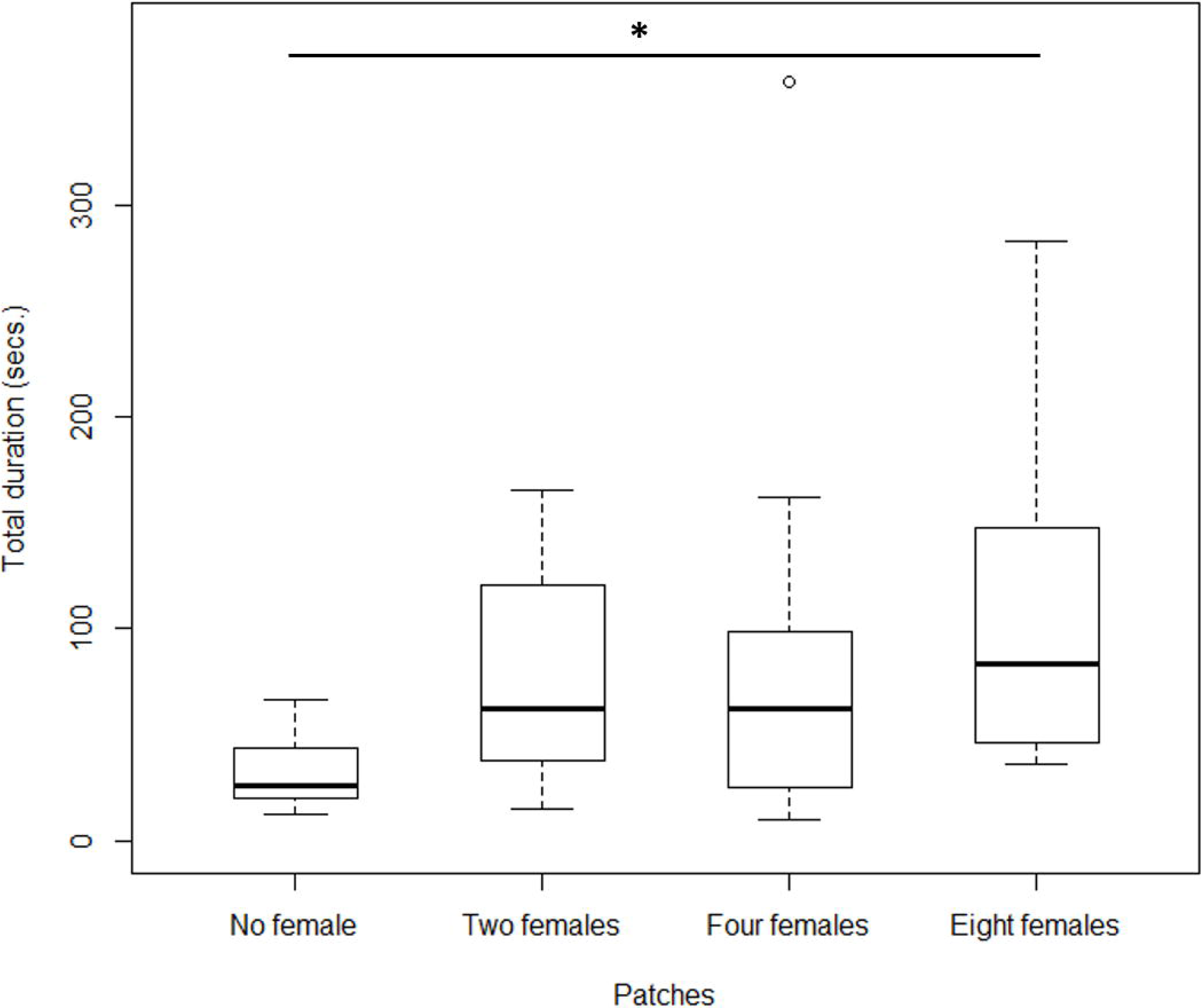
Boxplots showing the total duration (in secs.) spent the male in each of the chambers. Males spent significantly longer time in the chamber containing 2 females compared to the chamber with no females. The solid line with star indicates statistical significance between the two plots (p<0.05). The lower and upper ends of the box plots represent the 1st and 3rd quartiles, the horizontal line within each boxplot is the median, and the upper and lower whiskers are the 1.5× the interquartile range. Outlines are shown as open circles.

**Fig 3c.**
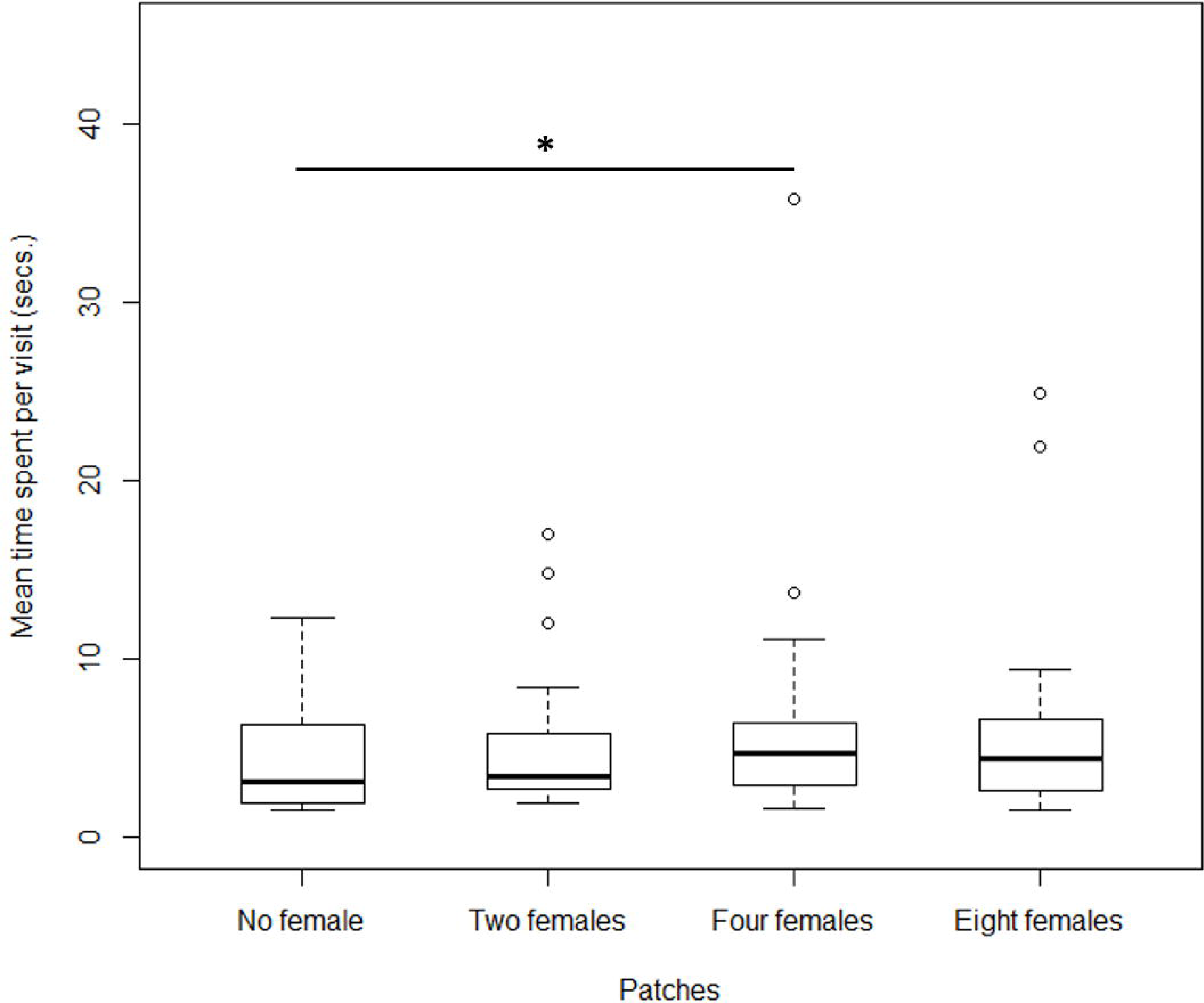
Boxplots showing the mean duration of each visit spent by the test male in each of the chambers. Males spent significantly longer time during every visit in the chamber containing 2 females compared to the chamber with no females. The solid line with star indicates statistical significance between the two plots (p<0.05). The lower and upper ends of the box plots represent the 1st and 3rd quartiles, the horizontal line within each boxplot is the median, and the upper and lower whiskers are the 1.5× the interquartile range. Outlines are shown as open circles.

### Association preference experiment with varying female and vegetation densities

#### For E1 set

Total number of visits was found to best fit a negative binomial distribution (AIC=441.75) and the model was constructed using ‘glmer.nb’ as mentioned above. The model (Wald type II χ2=34.95, df=3, p<0.001) revealed a significant effect of patch on total number of visits. (Table 2a). Post hoc tests showed significant difference in the total number of visits between H-L (V=238.5, p=0.002), H-N (V=204, p=0.002), M-L (V=247.5, p≤0.001) and M-N (V=256, p≤0.001) patches (Fig. 4a).

**Fig 4a.**
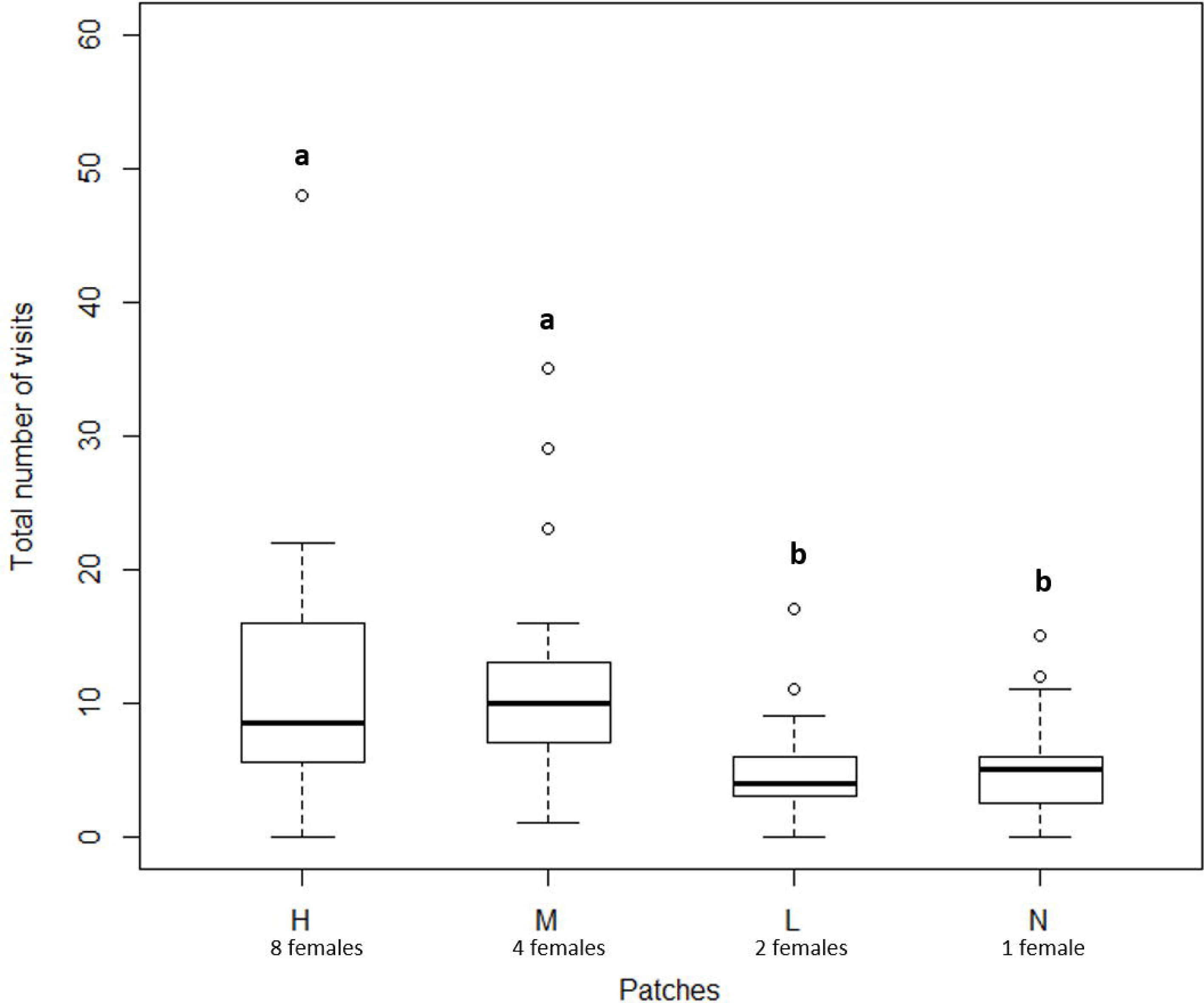
Boxplots showing the total number of visits by the male to the various chambers containing the females in varying density in vegetation presence (E1 set). High and medium chambers having eight females (and high plant density) and four females (and medium plant density) respectively had a higher number of visits by the male compared to the low (two females and low plant density) and null (one female and no plants) chambers. Similar alphabets placed above the plots indicate no statistical difference whereas dissimilar alphabets indicate significant statistical difference (p<0.05). The lower and upper ends of the box plots represent the 1st and 3rd quartiles, the horizontal line within each boxplot is the median, and the upper and lower whiskers are the 1.5× the interquartile range. Outlines are shown as open circles.

**Table 2.**
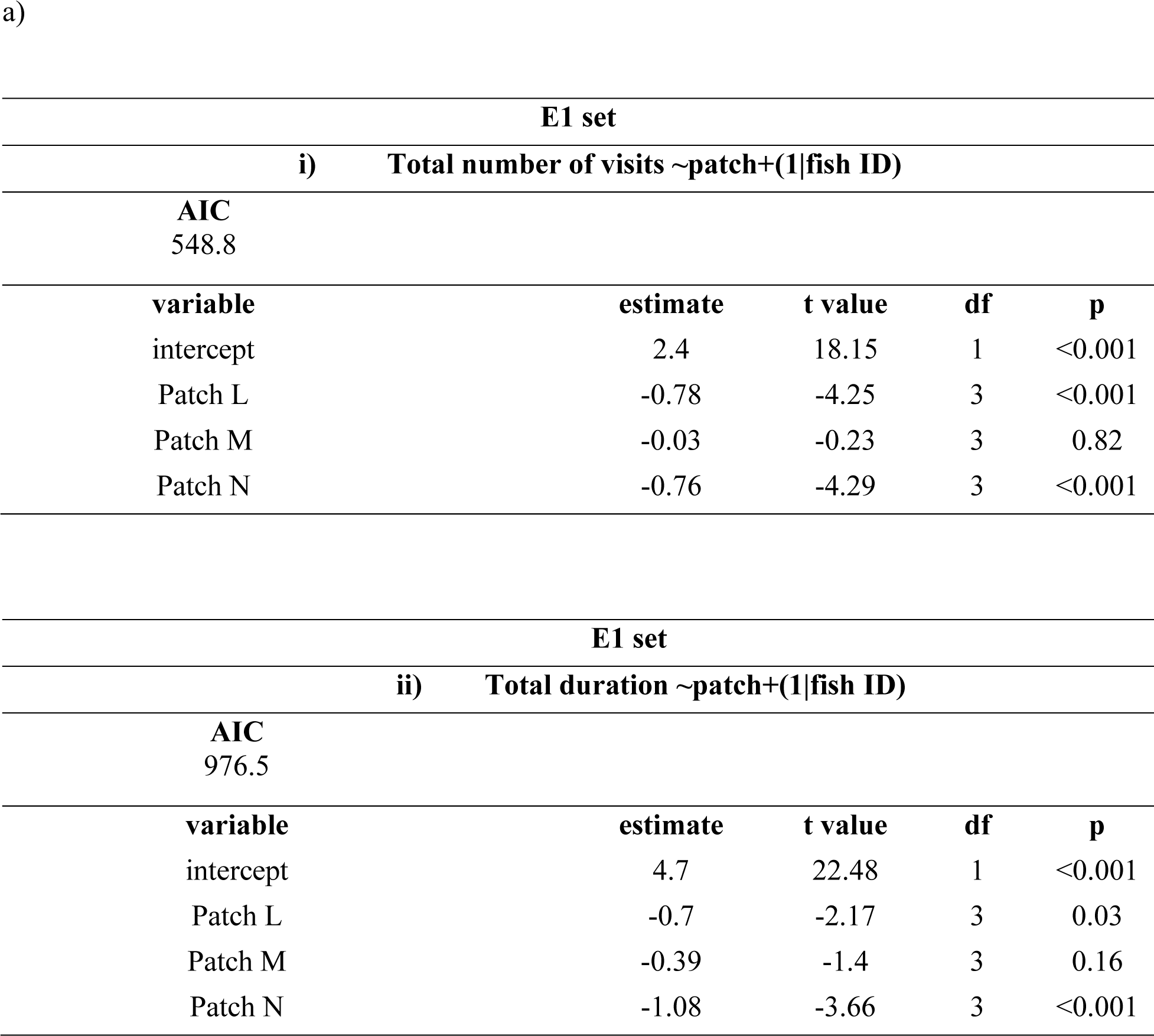
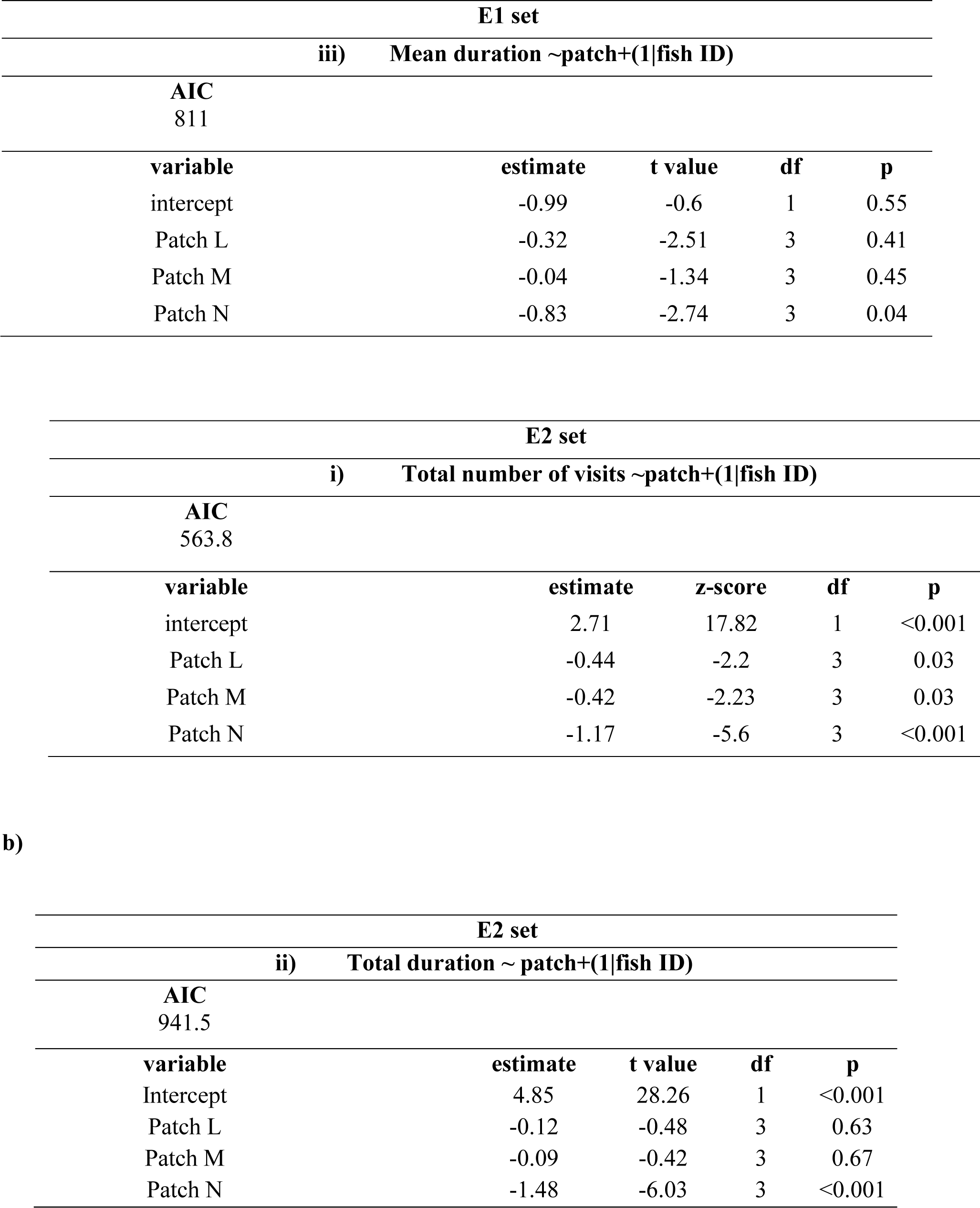
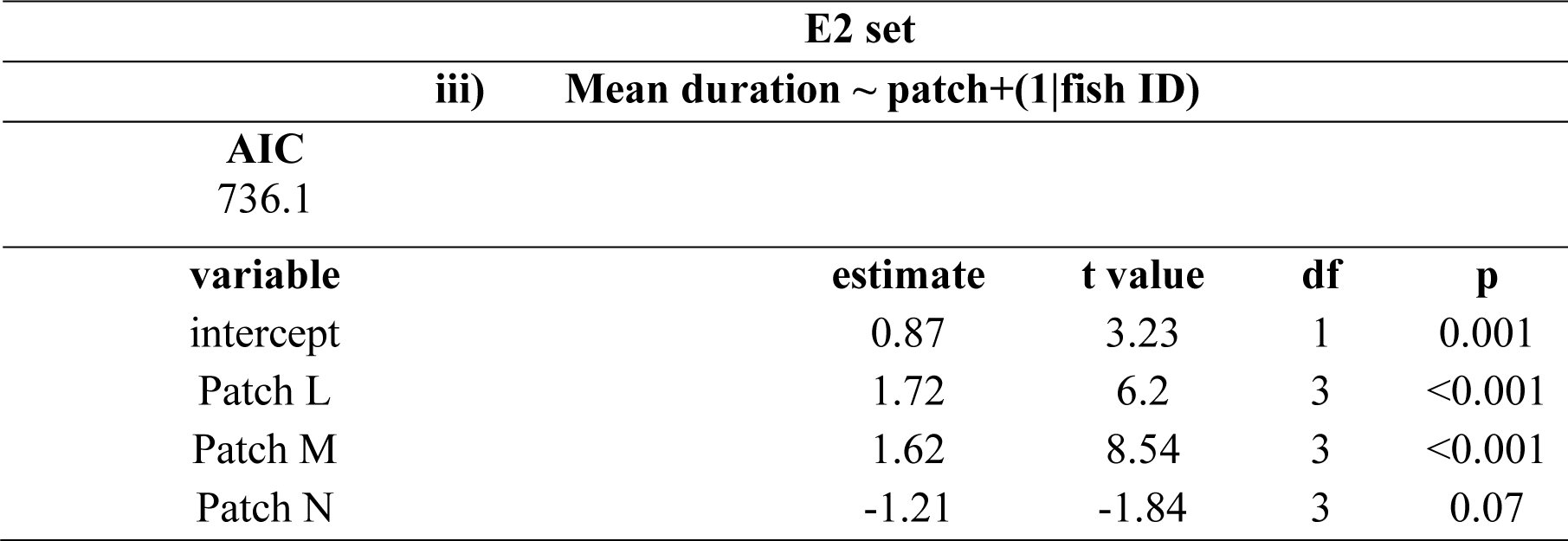
Summary of results of generalized linear mixed models indicating effect of patch (female number), tree (vegetation density) and fish id (random factor) on i) total number of visits, ii) total duration, iii) Mean duration of time spent at each patch. a) E1 set represents the experiments where female density in each patch increased with associated vegetation density. b) E2 set represents experiments where female density in patches decreases with vegetation density, except in the ‘no fish’ patch which had one plant near it. The AIC values of selected models along with estimate values, t-scores, degrees of freedom (df) and p values for each factor are shown. p values≤0.05 is significant. Patch2F, Patch4F, and Patch8F signify patches with 2, 4, and 8 females, respectively.

| E1 set |  |  |  |  |
| --- | --- | --- | --- | --- |
| i) Total number of visits ∼patch+(1\|fish ID) | | | | |
| AIC<br>548.8 |  |  |  |  |
| variable | estimate | t value | df | p |
| intercept | 2.4 | 18.15 | 1 | <0.001 |
| Patch L | -0.78 | -4.25 | 3 | <0.001 |
| Patch M | -0.03 | -0.23 | 3 | 0.82 |
| Patch N | -0.76 | -4.29 | 3 | <0.001 |
| ii) Total duration ∼patch+(1\|fish ID) | | | | |
| AIC<br>976.5 |  |  |  |  |
| variable | estimate | t value | df | p |
| intercept | 4.7 | 22.48 | 1 | <0.001 |
| Patch L | -0.7 | -2.17 | 3 | 0.03 |
| Patch M | -0.39 | -1.4 | 3 | 0.16 |
| Patch N | -1.08 | -3.66 | 3 | <0.001 |
| iii) Mean duration ~patch+(1\|fish ID) |  |  |  |  |
| AIC<br>811 |  |  |  |  |
| variable | estimate | t value | df | p |
| intercept | -0.99 | -0.6 | 1 | 0.55 |
| Patch L | -0.32 | -2.51 | 3 | 0.41 |
| Patch M | -0.04 | -1.34 | 3 | 0.45 |
| Patch N | -0.83 | -2.74 | 3 | 0.04 |

| E2 set |  |  |  |  |
| --- | --- | --- | --- | --- |
| i) Total number of visits ~patch+(1\|fish ID) |  |  |  |  |
| AIC<br>563.8 |  |  |  |  |
| variable | estimate | z-score | df | p |
| intercept | 2.71 | 17.82 | 1 | <0.001 |
| Patch L | -0.44 | -2.2 | 3 | 0.03 |
| Patch M | -0.42 | -2.23 | 3 | 0.03 |
| Patch N | -1.17 | -5.6 | 3 | <0.001 |

b)
| E2 set |  |  |  |  |
| --- | --- | --- | --- | --- |
| ii) Total duration ~ patch+(1\|fish ID) |  |  |  |  |
| AIC<br>941.5 |  |  |  |  |
| variable | estimate | t value | df | p |
| Intercept | 4.85 | 28.26 | 1 | <0.001 |
| Patch L | -0.12 | -0.48 | 3 | 0.63 |
| Patch M | -0.09 | -0.42 | 3 | 0.67 |
| Patch N | -1.48 | -6.03 | 3 | <0.001 |

| E2 set |  |  |  |  |
| --- | --- | --- | --- | --- |
| iii) Mean duration ~ patch+(1\|fish ID) |  |  |  |  |
| AIC |  |  |  |  |
| 736.1 |  |  |  |  |
| variable | estimate | t value | df | p |
| intercept | 0.87 | 3.23 | 1 | 0.001 |
| Patch L | 1.72 | 6.2 | 3 | <0.001 |
| Patch M | 1.62 | 8.54 | 3 | <0.001 |
| Patch N | -1.21 | -1.84 | 3 | 0.07 |

GLMM for total duration (Wald type II χ2=13.95, df=3, p=0.002) revealed a significant effect of patches (Table 2a). Post hoc paired Wilcoxon tests between the groups showed significant effects between H-N (V=233, p=0.004) and M-N (V=247, p=0.004) patches (Fig. 4b).

**Fig 4b.**
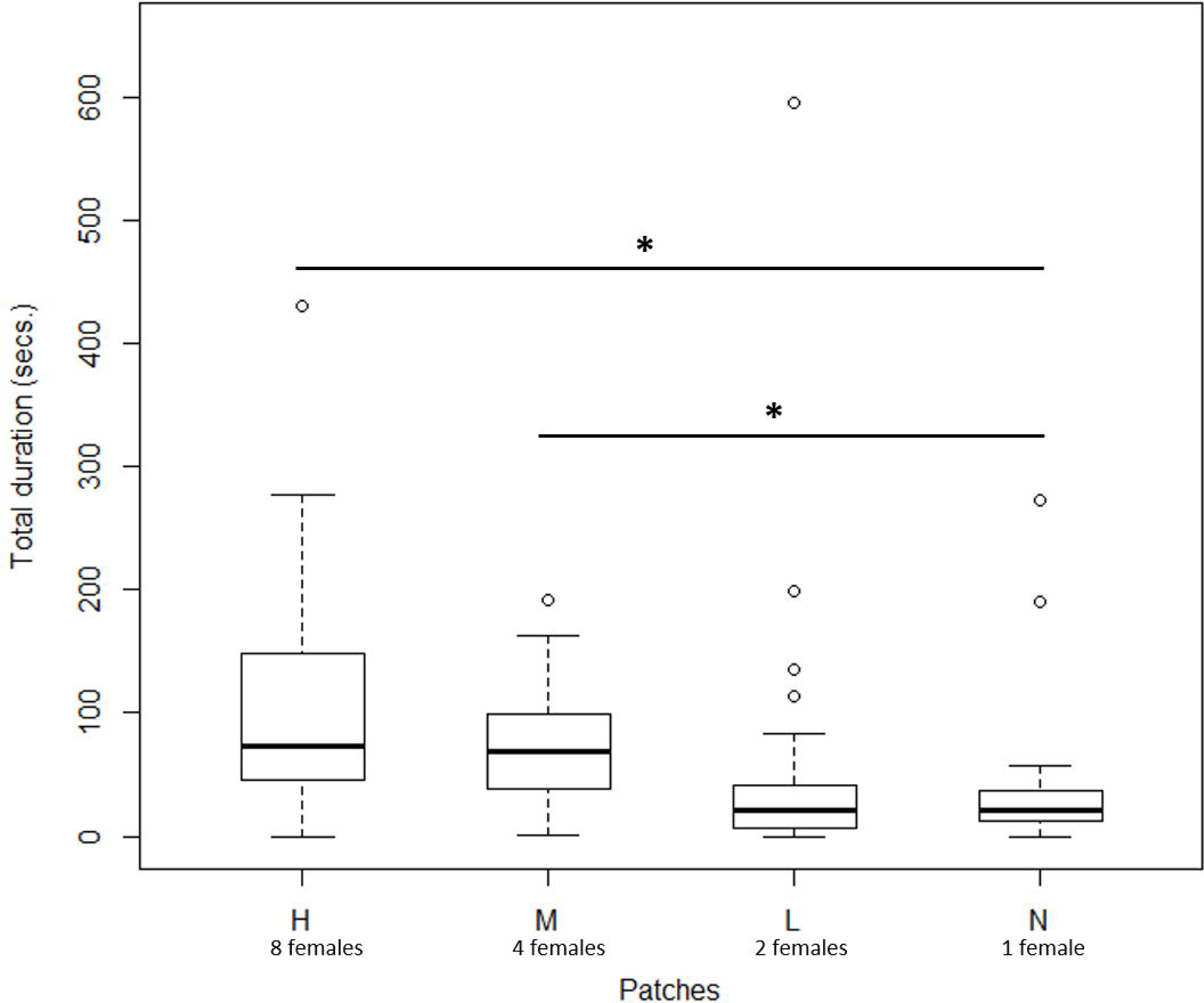
Boxplots showing the total duration (in secs.) spent the male in each of the chamber (E1 set). Males spent a significantly longer time in the high and medium chamber compared to the null chamber. The solid line with star indicates statistical significance between the two plots (p<0.05). The lower and upper ends of the box plots represent the 1st and 3rd quartiles, the horizontal line within each boxplot is the median, and the upper and lower whiskers are the 1.5× the interquartile range. Outlines are shown as open circles.

For the model for mean duration of time, patch was a significant factor (Wald type II χ2=14.31, df=3, p=0.002). Post hoc Wilcoxon tests showed a significant difference between H-N (V= 220, p=0.01) and M-N (V= 224, p=0.03) (Fig. 4c) (Table 2a).

**Fig 4c.**
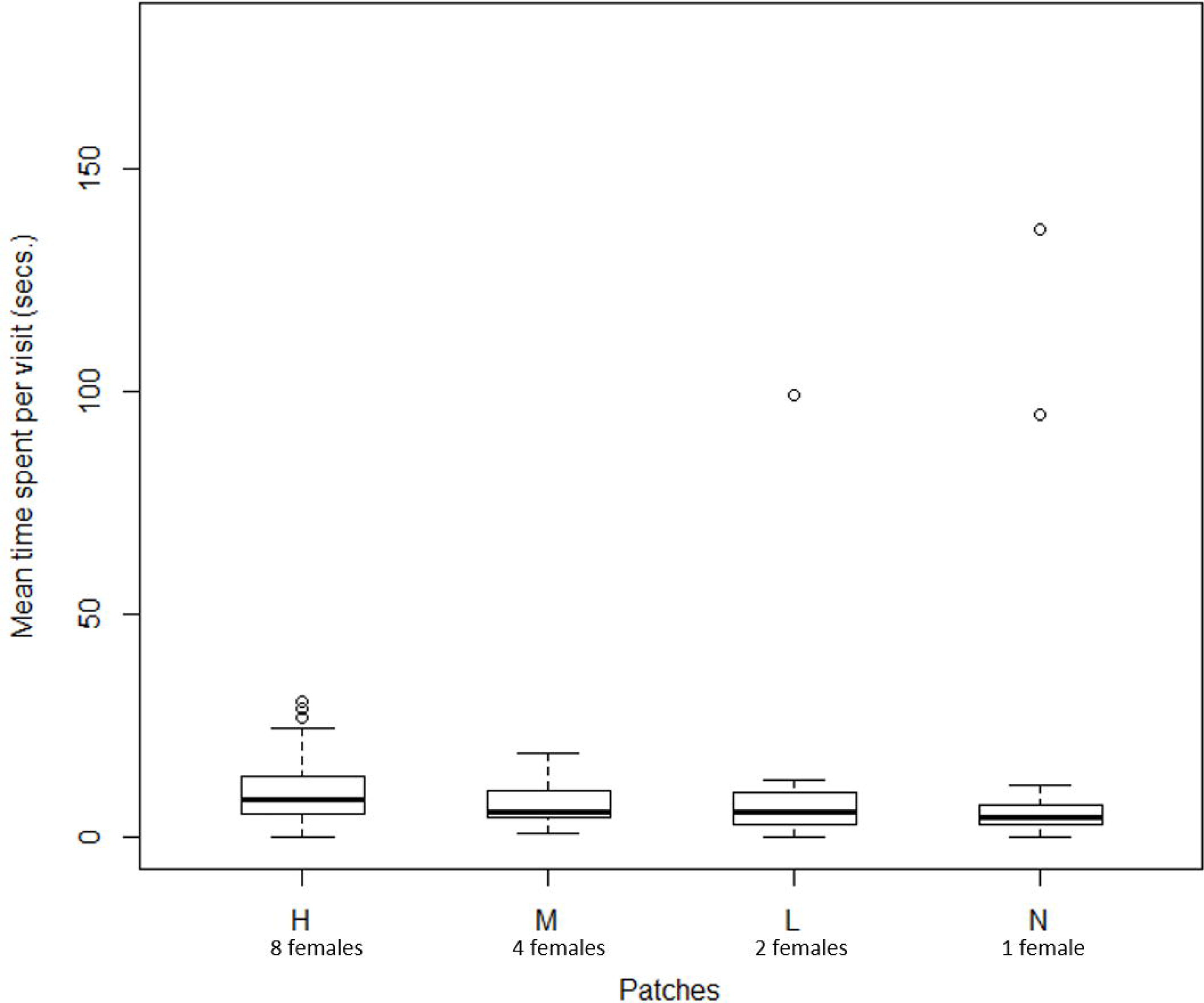
Boxplots showing the mean duration of each visit spent by the test male in each of the chamber (E1 set). Males spent statistically similar times in all the four chambers. The lower and upper ends of the box plots represent the 1st and 3rd quartiles, the horizontal line within each boxplot is the median, and the upper and lower whiskers are the 1.5× the interquartile range. Outlines are shown as open circles.

#### For E2 set

Negative binomial glmm for the total number of visits showed patches (Wald type II χ2=31.53, df=3, p<0.0010.002) as a significant factor (Table 2b). Paired comparisons show significant differences between H-L (V=256, p=0.002), H-N (V=278.5, p<0.001), and M-N (V=214.5, p<0.001) (Fig. 5a).

**Fig 5a.**
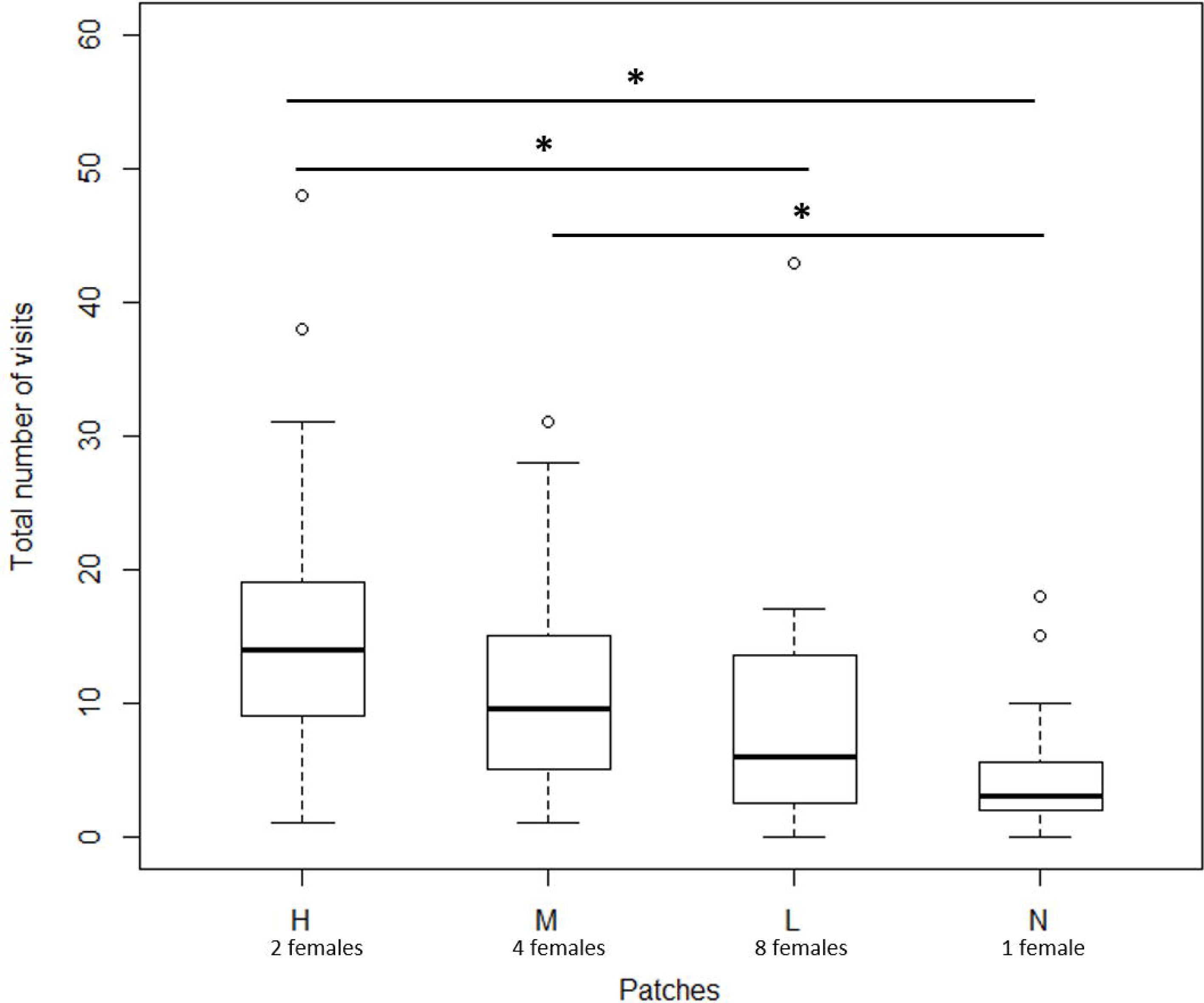
Boxplots showing the total number of visits by the male to the various chambers (E2 set). Males visited the high chamber (2 females and high plant density) significantly more than the low (eight females with low plant density) or null (one female with no plants) chambers. Medium (four females with medium plant density) chamber also received a significantly higher number of visits than the null chamber. The solid line with star indicates statistical significance between the two plots (p<0.05). The lower and upper ends of the box plots represent the 1st and 3rd quartiles, the horizontal line within each boxplot is the median, and the upper and lower whiskers are the 1.5× the interquartile range. Outlines are shown as open circles.

Like E1, log=link function was used to analyze the total duration (“Gamma” family) and the mean time per visit (“Gaussian” family) data. Patch significantly affected total time spent in a patch (Wald type II χ2=48.94, df=3, p<0.001) (Table 2b). Post hoc Wilcoxon shows a significant difference between H-N (V=278, p<0.001), M-N (V=293, p<0.001) and L-N (V=224, p0.002) (Fig. 5b).

**Fig 5b.**
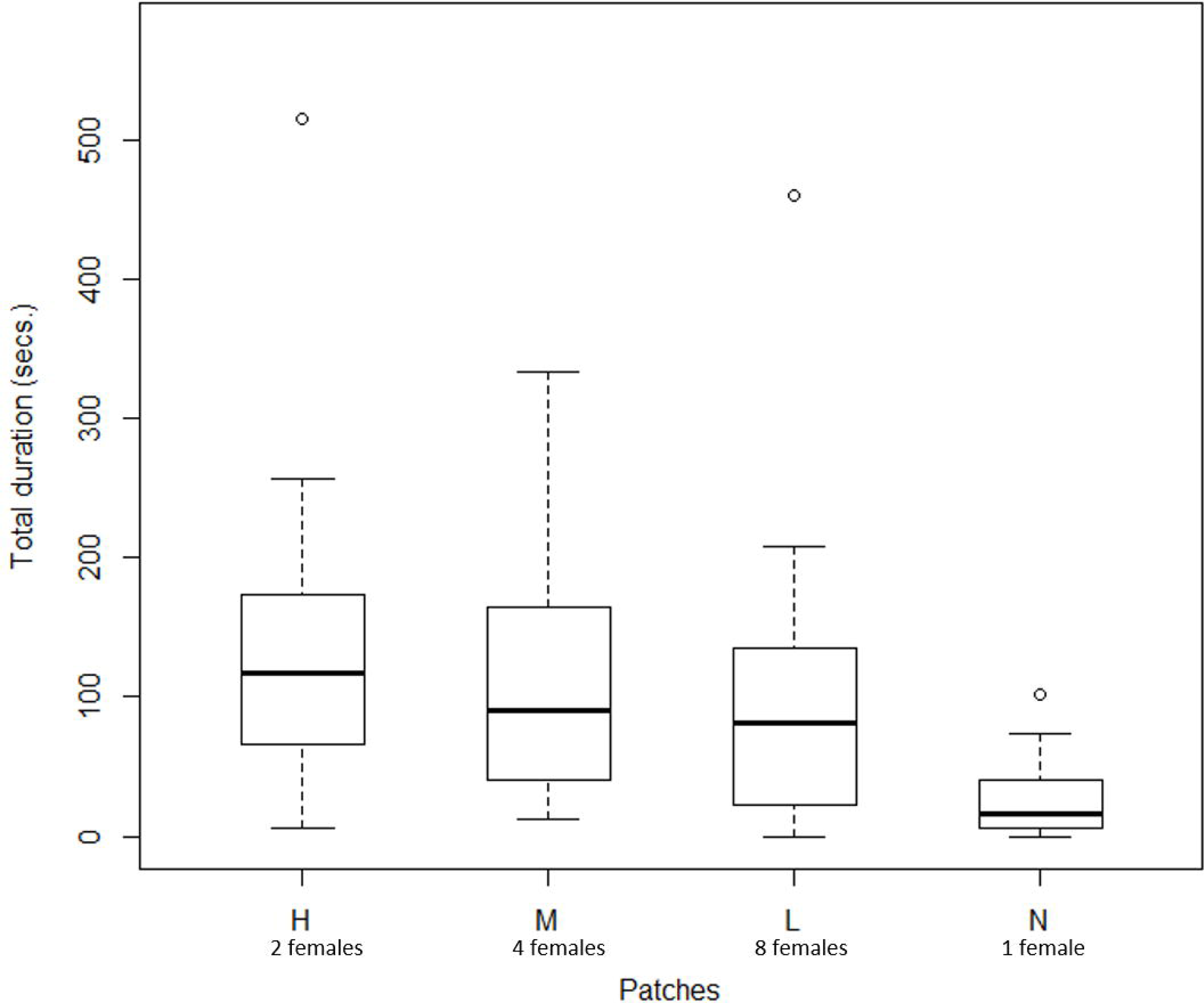
Boxplots showing the total duration (in secs.) spent the male in each of the chamber (E2 set). The test males spent statistically similar times in all the four patches. The lower and upper ends of the box plots represent the 1st and 3rd quartiles, the horizontal line within each boxplot is the median, and the upper and lower whiskers are the 1.5× the interquartile range. Outlines are shown as open circles.

The mean time per visit was significantly affected by patches (Wald type II χ2=90.23, df=3, p<0.001) (Table 2b). Posthoc paired tests revealed a significant difference only in M-N (V=251, p=0.003) (Fig. 5c).

**Fig 5c.**
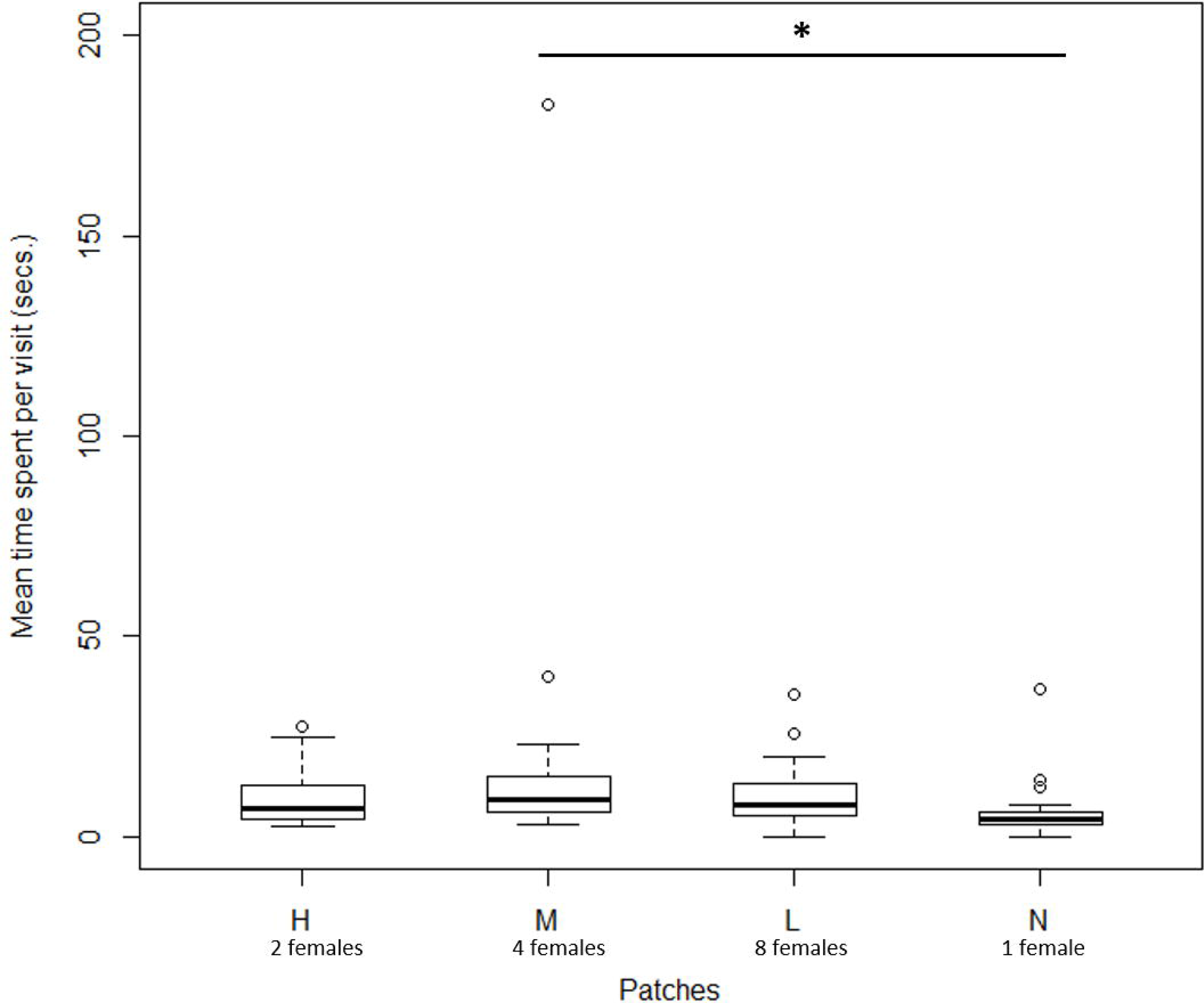
Boxplots showing the mean duration of each visit spent by the test male in each of the chamber (E1 set). The duration mean was significantly higher in the medium (M) patch compared to the null patch. The solid line with star indicates statistical significance between the two plots (p<0.05). The lower and upper ends of the box plots represent the 1st and 3rd quartiles, the horizontal line within each boxplot is the median, and the upper and lower whiskers are the 1.5× the interquartile range. Outlines are shown as open circles.

We additionally performed pair-wise Wilcoxon tests between H and L patches of E1 and E2 sets to compare the total number of visits and total duration spent across our two experimental setups. Total number of visits did not differ for H patch of E1 and E2 (v=79.5, p=0.08) or L patch between E1 and E2 (v=87.5, p=0.13). Total duration was also comparable for H patches across the two sets (v=120, p=0.4) along with L patches (v=88, p=0.79).

## DISCUSSION

Our study aimed to understand the influence of the density of females (potential mates) in determining the association preference of males with a particular patch. Furthermore, we also investigated if vegetation cover associated with the female patches influenced male association preferences. Our results revealed that while there was no clear effect of female number on the males’ association preferences, we found that male association is significantly greater when there was a higher vegetation cover, irrespective of female number The findings are contrary to the general expectation that male association preferences in the context of mate choice would be based primarily on female numbers. In natural habitats, however, male choices would be based on balancing of the costs and benefits from more than one ecological factor present in their immediate surroundings (Sih and Krupa 1995; Wong and Jennions 2003). We discuss the results in detail in the following sections.

### Effect of female number on association preferences

When presented a multichoice setup with varying female numbers, we found that males clearly prefer a patch with females as all the three patches that housed females had a significantly higher number of visits than the null or zero-female containing patch. However, within the three female-containing patches, we did not see any significant difference in the total number of visits or total time spent. The lack of as clear choice for the high female numbers patch could be a result of the lack of sharp difference between the numbers. Earlier literature indicates that zebrafish are unable to distinguish between group sizes when the number of individuals in the choice groups are not sharply different (Potrich et al 2015; Seguin and Gerlai 2017). The physical state and receptivity of females (McNamara et al. 2004) could also influence decision making in males (Byrne and Rice 2006). This effect will be all the more pronounced in lower female densities and as in higher densities, the sheer greater number might be a strong enough stimulus for the male. Furthermore, other factors such as predation pressure or risk could also play a role in differential preference for patches (Seghers 1981; Hagar and Helfman 1991). Thus, schooling tendency itself could play a role in determining preference for patches varying in individual densities. Absence of a preferential choice for associating with patches varying in female numbers in our study could be due to there being no ‘threat’ or survival cost associated with a specific patch. Further experimental investigations are warranted to understand other underlying factors that can determine male zebrafish responses to changes in female density.

### Association preferences in the presence of varying vegetation density

Our results revealed that the presence of vegetation in patches significantly affected male association preferences. In E1, where female number was proportional to tree density, we found that high (H) and medium (M) density patches were preferred in all measured parameters of association preference i.e., total number of visits, total time spent and average time spent per visit. However, there is a decrease in the number of visits as well as total time spent in low (L) and null (N) patches. In E2, our H patch was actually associated with only two females and L patch was associated with eight females. However, despite fewer number of females, the pattern of male association stayed the same with H still getting significantly higher number of visits than the L patch. We, however, did find that the males spent comparable amounts of time in the H, M and L patches all three of which were significantly higher than the N patch. Though they visited the L patch less often they did spend greater time per visit due to the high number of females in that patch. This indicates a probable interaction between the two factors vegetation density and female number influencing the male’s association patterns. Further analysis between the two experimental groups found no difference in any of the parameters between the H patches in E1 and E2 conditions. So, although, there were both high and low numbers of females associated with H patch in these conditions, the male visit was not affected which indicates the possible sheltering role the high vegetation density plays in the H patch. Similarly, no difference was observed between the L patches in E1 and E2 conditions. Even though L patch had very minimal vegetation in E2 allowing easier perception of higher number of females in E2, it still did not change the association patterns. These results, together point towards a more important role of vegetation cover than we had expected. Spence et al. (2007) studied the role of vegetation and oviposition site quality in egg-laying decisions by female zebrafish. They found that females prefer sites with vegetation cover and this might reflect greater survival of the larvae. In the wild, zebrafish are generally occurred closer to vegetation cover around the edge of water bodies and furthermore, prefer to spawn in such areas (Spence et al. 2006). Vegetation allows for the larvae to attach to them till the swim bladder is inflated (Laale 1977) thus increasing their survivability. Kistler et al. (2011) showed that complex structured environments are preferred by even laboratory-bred strains of zebrafish and checker barbs. They speculate that this preference for vegetation cover is a predator-avoidance trait, despite lab-reared strains experiencing relaxed selection for anti-predator responses, indicating an intrinsic affinity for vegetation cover. The importance of vegetation cover has also been studied as an enrichment factor in zebrafish housing tanks in laboratories (Wilkes et al. 2012; Keck et al. 2015). Our findings underline the additional effects of vegetation in important behaviors such as mate search and mate association among zebrafish.

Male mate search dynamics are one of the most widely studied aspects of reproductive ecology. There are several theoretical hypotheses that delineate search algorithms for the searcher (Real 1991; Johnstone 1997) which have also been tested empirically (Fiske and Kålås 1995; Murphy 2012; Kozak et al. 2013; Deb and Balakrishnan 2014). Zebrafish is a sexually monomorphic species and are only reported to show coloration changes during courtship ephemerally (Hutter et al. 2012). In such species, demographic and ecological factors would strongly influence mating strategies. Our current lacuna in understanding various aspects of mating strategies in fish makes the zebrafish a useful model organism to ask questions about ecological factors controlling mate choice and mating behavior.

To our knowledge, our study is the first of its kind that investigates the mate-search paradigm in zebrafish males in the context of female number along with habitat complexity. The use of the multi-choice paradigm with a gradient in stimuli intensity represents a more realistic approach to understand complex behaviors such as mate search. The dimensions of the experimental arena allowed the male to sequentially visit each patch and sample it. This also impacted the cost of traveling between patches and a cost associated with sampling each patch having multiple potential mates. Even though it is an artificial design it was necessary to establish individual behavioral preferences that can allow us a deeper understanding of how intra-group dynamics might emerge. Our results point towards a complex interaction between female number and vegetation in influencing male association. These findings also indicate the need for a deeper understanding of how aquatic vegetation modulates mating strategies, to help provide informed management plans through incorporation of native aquatic foliage for the conservation of endangered or threatened aquatic species.

## Ethical Statement

The study complied with the existing rules and guidelines outlined by the Committee for the Purpose of Control and Supervision of Experiments on Animals (CPCSEA), Government of India, the Institutional Animal Ethics Committee’s (IAEC) and guidelines of Indian Institute of Science Education and Research (IISER) Kolkata. No animals were euthanized or sacrificed during any part of the study, and behavioral observations were conducted without any chemical treatment on the individuals. At the end of the experiments, all fish were returned to stock tanks and continued to be maintained in the laboratory.

## Acknowledgments

The authors would like to thank the Indian Institute of Science Education and Research (IISER)-Kolkata (India) for providing infrastructural and financial support for the study. AG received Junior and Senior Research Fellowships from University Grants Commission (UGC), Government of India. Help from Rubina Mondal and Danita Daniel for routine maintenance and upkeep of laboratory and fish tanks is deeply appreciated.

## Bibliography

Bates D, Maechler M, Bolker B, Walker S (2015) Fitting Linear Mixed-Effects Models Using lme4. JStat Soft, 67(1): 1–48.

Berglund A (1993) Risky sex: male pipefishes mate at random in the presence of a predator. Anim Behav. 46(1): 169–175.

Bierbach D, Stadler S, Jung CT, Kunkel B, Ziege M, Plath M, Schulte M, Herrmann N, Tobler M, Riesch R, Klaus S, Indy JR, Arias-Rodriguez L (2011) Predator-induced changes in female mating preferences: Innate and experiential effects. BMC Evol Biol. 11(1): 190. doi.org/10.1186/1471-2148-11-190

Byrne PG, Rice WR (2006) Evidence for adaptive male mate choice in the fruit fly *Drosophila melanogaster*. Proc R Soc B Biol Sci. 273: 917–922. doi.org/10.1098/rspb.2005.3372

Carfagnini AG, Rodd FH, Jeffers KB, Bruce AEE (2009) The effects of habitat complexity on aggression and fecundity in zebrafish (*Danio rerio*). Environ Biol Fishes 86: 403–409. doi.org/10.1007/s10641-009-9539-7

Christensen, B., & Persson, L. (1993). Species-specific antipredatory behaviours: effects on prey choice in different habitats. Behavioral Ecology and Sociobiology, 32(1), 1–9.

Darrow, K.O., Harris, W. A (2004) Characterization and development of courtship in zebrafish, *Danio rerio*. Zebrafish 1, 40–45. doi.org/10.1089/154585404774101662

Deb R, Balakrishnan R (2014) The opportunity for sampling: The ecological context of female mate choice. Behav Ecol. 25: 967–974. doi.org/10.1093/beheco/aru072

Delaney M, Follet C, Ryan N, Hanney N, Lusk-Yablick J, Gerlach G (2002) Social interaction and distribution of female zebrafish (*Danio rerio*) in a large aquarium. Biol Bull. 203(2): 240–241.

Dugatkin LA, FitzGerald GJ (1997) Sexual selection. In: Behavioural Ecology of Teleost Fishes (Godin, JJ ed.). Oxford Univ. Press, New York, pp. 266–284.

Edward DA, Chapman T. 2011. The evolution and significance of male mate choice. Tr Ecol Evol. 26: 647–654. doi.org/10.1016/j.tree.2011.07.012

Eens M, Pinxten R (2000) Sex-role reversal in vertebrates: Behavioural and endocrinological accounts. Behav Proc. 51: 135–147. doi.org/10.1016/S0376-6357(00)00124-8

Fiske P, Kålås JA (1995) Mate sampling and copulation behaviour of great snipe females. Anim Behav 49(1): 209–219.

Forsgren E (1992) Predation Risk Affects Mate Choice in a Gobiid Fish. Am Nat. 140 (6): 1041–1049. doi.org/10.1086/285455

Friard O, Gamba M (2016) BORIS: a free, versatile open source event logging software for video/audio coding and live observations. Meth Ecol. Evol. 7(11): 1325–1330.

Gerlach G (2006) Pheromonal regulation of reproductive success in female zebrafish: female suppression and male enhancement. Anim Behav. 72: 1119–1124. doi.org/10.1016/j.anbehav.2006.03.009

Gwynne DT (2016) Sexual Selection: Roles Evolving. Curr Biol. 26: R935–R936. doi.org/10.1016/j.cub.2016.08.063

Hagar MC, Helfman GS (1991) Safety in numbers: shoal size choice by minnows under predation threat. Behav Ecol Sociobiol 29:271–276

Hothorn T, Bretz F, Westfall P (2008) Simultaneous Inference in General Parametric Models. Biomet J. 50(3): 346–363.

Hutter S, Hettyey A, Penn DJ, Zala SM (2012) Ephemeral Sexual Dichromatism in Zebrafish (*Danio rerio*). Ethology 118: 1208–1218. doi.org/10.1111/eth.12027

Jirotkul M (1999) Population density influences male-male competition in guppies. Anim. Behav. 58(6): 1169–1175. doi.org/10.1006/anbe.1999.1248

Johnstone RA (1997) The tactics of mutual mate choice and competitive search. Behav Ecol Sociobiol. 40(1): 51–59.

Keck VA, Swift LL, Boyd KL, Edgerton DS, Hajizadeh S, Dupont WD, Lawrence C (2015) Effects of habitat complexity on pair-housed zebrafish. J Am Assoc Lab Anim Sci. 54: 378–383.

Kistler C, Hegglin D, Würbel H, König B (2011) Preference for structured environment in zebrafish (*Danio rerio*) and checker barbs (*Puntius oligolepis*). Appl Anim Behav Sci. 135: 318–327. doi.org/10.1016/j.applanim.2011.10.014

Kozak GM, Head ML, Lackey ACR, Boughman, JW (2013) Sequential mate choice and sexual isolation in threespine stickleback species. J Evol Biol. 26: 130–140. doi.org/10.1111/jeb.12034

Kvarnemo C, Ahnesjö I (1996) The dynamics of operational sex ratios and competition for mates. T Ecol Evol 11: 404–408. doi.org/10.1016/0169-5347(96)10056-2

Laale HW (1977) The biology and use of zebrafish, *Brachydanio rerio* in fisheries research. A literature review. J Fish Biol. 10(2): 121–173. doi.org/10.1111/j.1095-8649.1977.tb04049.x

McNamara KB, Jones TM, Elgar MA (2004) Female reproductive status and mate choice in the hide beetle, *Dermestes maculatus*. J Insect Behav. 17: 337–352. doi.org/10.1023/B:JOIR.0000031535.00373.b1

Murphy CG (2012) Simultaneous mate-sampling by female barking treefrogs (*Hyla gratiosa*). Behav Ecol. 23: 1162–1169. doi.org/10.1093/beheco/ars093

Nasiadka A, Clark MD (2012) Zebrafish breeding in the laboratory environment. ILAR J. 53: 161–168. doi.org/10.1093/ilar.53.2.161

Ostrand KG, Braeutigam BJ, Wahl DH (2004) Consequences of Vegetation Density and Prey Species on Spotted Gar Foraging. Trans. Am Fish Soc. 133: 794–800. doi.org/10.1577/t03-111.1

Owen MA, Rohrer K, Howard RD (2012) Mate choice for a novel male phenotype in zebrafish, *Danio rerio*. Anim Behav. 83: 811–820. doi.org/10.1016/j.anbehav.2011.12.029

Potrich, D., Sovrano, V. A., Stancher, G., & Vallortigara, G. (2015). Quantity discrimination by zebrafish (Danio rerio). Journal of Comparative Psychology, 129(4), 388.

Pyron M (2003) Female preferences and male-male interactions in zebrafish (*Danio rerio*). Can J Zool. 81: 122–125. doi.org/10.1139/z02-229

Real LA (1991) Search theory and mate choice. II. Mutual interaction, assortative mating, and equilibrium variation in male and female fitness. Am Nat. 138(4): 901–917.

Roy T, Bhat A (2015) Can outcomes of dyadic interactions be consistent across contexts among wild zebrafish? R Soc Open Sci. 2 (11): 150282. doi.org/10.1098/rsos.150282

Schroeder P, Jones S, Young IS, Sneddon LU (2014) What do zebrafish want? Impact of social grouping dominance and gender on preference for enrichment. Lab Anim. 48: 328–337. doi.org/10.1177/0023677214538239

Seghers BH (1981) Facultative schooling behaviour in the spottail shiner (*Notropis hudsonius*): possible costs and benefits. Environ Biol Fish 6:21–24

Seguin, D., & Gerlai, R. (2017). Zebrafish prefer larger to smaller shoals: analysis of quantity estimation in a genetically tractable model organism. Animal cognition, 20(5), 813–821.

Sih A, Krupa JJ (1995) Interacting effects of predation risk and male and female density on male/female conflicts and mating dynamics of stream water striders. Behav Ecol. 6(3): 316–325.

Skinner AMJ, Watt PJ (2007) Strategic egg allocation in the zebrafish, *Danio rerio*. Behav Ecol. 18: 905–909. doi.org/10.1093/beheco/arm059

Spence R, Smith C (2006) Mating preference of female zebrafish, *Danio rerio*, in relation to male dominance. Behav. Ecol. 17: 779–783. doi.org/10.1093/beheco/arl016

Spence R, Ashton R, Smith C (2007) Oviposition decisions are mediated by spawning site quality in wild and domesticated zebrafish, *Danio rerio*. Behaviour 144: 953–966. doi.org/10.1163/156853907781492726

Trivers RL (1972) Parental investment and sexual selection. In ‘Sexual Selection and the Descent of Man’.(Ed. B. Campbell.) pp. 136–179. Aldinc: Chicago.

Uusi-Heikkilä S, Böckenhoff L, Wolter C, Arlinghaus R (2012) Differential allocation by female zebrafish (*Danio rerio*) to different-sized males--an example in a fish species lacking parental care. PLoS One 7: e48317. doi.org/10.1371/journal.pone.0048317

Wagner W (1998) Measuring female mating preferences. Anim Behav. 55: 1029–42. doi.org/10.1006/anbe.1997.0635

Weir LK, Grant JWA, Hutchings, JA (2011) The influence of operational sex ratio on the intensity of competition for mates. Am Nat. 177: 167–176. doi.org/10.1086/657918

Weiss SL, Dubin M (2018) Male mate choice as differential investment in contest competition is affected by female ornament expression. Curr Zool. 64: 335–344. doi.org/10.1093/cz/zoy023

Wilkes L, Owen SF, Readman GD, Sloman KA, Wilson RW (2012) Does structural enrichment for toxicology studies improve zebrafish welfare? Appl Anim Behav Sci. 139: 143–150. doi.org/10.1016/j.applanim.2012.03.011

Willis PM, Ryan MJ, Rosenthal GG (2011) Encounter rates with conspecific males influence female mate choice in a naturally hybridizing fish. Behav Ecol. 22: 1234–1240. doi.org/10.1093/beheco/arr119

Wong BBM, Jennions MD (2003) Costs influence male mate choice in a freshwater fish. Proc. R. Soc. London. (Ser. B) Biol. Sci. 270: S36–S38. doi.org/10.1098/rsbl.2003.0003

